# Hepatic Nrf1 (*Nfe2l1*) promotes VLDL dependent liver defense against sepsis

**DOI:** 10.1101/2024.07.04.602118

**Authors:** Michael J. Trites, Lei Li, May G. Akl, Aidan Hydomako, Scott B. Widenmaier

**Affiliations:** Department of Anatomy, Physiology, and Pharmacology, University of Saskatchewan, Saskatoon, SK, Canada

**Author notes:** Corresponding author information Scott B. Widenmaier, 107 Wiggins Road, Health Sciences Building Saskatoon, SK S7N 5E5, Canada.

**Keywords:** Nrf1, Nfe2l1, liver, hepatocyte, disease tolerance, triglyceride, sepsis, stress defense

## Abstract

Sepsis causes mortality by triggering organ damage. Interest has emerged in stimulating disease tolerance to reduce organ damage. Liver plays a role in disease tolerance by mediating defensive adaptations, but sepsis-induced liver damage limit these effects. Here, we investigated whether stress defending transcription factors nuclear factor erythroid 2 related factor-1 (Nrf1) and -2 (Nrf2) in hepatocytes protect against sepsis. Using mice, we evaluated responses by hepatic Nrf1 and Nrf2 during sepsis triggered by lipopolysaccharide or *Escherichia coli*. We also genetically altered hepatic Nrf1 and Nrf2 activity to determine the protective role of these factors in sepsis. Our results show hepatic Nrf1 and Nrf2 activity is reduced in severe sepsis and hepatic Nrf1, but not Nrf2, deficiency predisposes for hypothermia and mortality. In contrast, enhancing hepatic Nrf1 activity protects against hypothermia and improves survival. Moreover, in sepsis hepatic Nrf1 deficiency reduces VLDL secretion whereas enhancing hepatic Nrf1 increases VLDL secretion, and inhibiting VLDL secretion with lomitapide obstructs protective actions of hepatic Nrf1. Gene expression profiles suggest Nrf1 promotes this effect by inducing stress defenses. Hence, we show mortality in sepsis may result from impaired stress defense and that hepatic Nrf1 improves disease tolerance during sepsis by promoting VLDL dependent liver defense.

## Introduction

Sepsis is a life-threatening organ dysfunction caused by a dysregulated response to infection (1). In hospital mortality is greater than 10 % for sepsis and 40 % for septic shock, and survivors have poor long-term prognosis (1–5). While current treatments focus on eliminating the infection (e.g., antibiotics) and hemodynamic stabilization (e.g., administer fluid and vasopressor), there is a recognized need for adjunct therapy that can mitigate organ damage and restore organ function to improve patient outcome (2, 6). During infection, the host has two defense strategies: resistance and tolerance (7, 8). Resistance is mediated by the immune system and involves inflammatory processes that can detect and eliminate the pathogen. However, this destructive and energetically costly process exerts collateral tissue damage that, in sepsis, leads to declining organ function and fitness (9). In contrast, tolerance is a strategy in which host defenses reduce susceptibility to tissue damage, irrespective of pathogen burden, resulting in preserved organ function and fitness (7–9). The mechanisms underlying disease tolerance and how they may be harnessed to improve the outcome of patients with sepsis is an emerging area of great interest (2, 6–10).

Metabolic adaptations that protect against organ damage underlie disease tolerance (9–13). In sepsis models using bacteria or endotoxin, reduced tissue damage and improved survival has been linked to adaptive shifts in glucose, ketone, heme, lipid, lipoprotein, and energy metabolism (14–23), which are similar to adaptive shifts during starvation and regulated by reactive oxygen species (ROS) and hormones such as glucocorticoid, fibroblast growth factor 21 (FGF21), growth and differentiation factor 15 (GDF15), and growth hormone (9, 13, 18, 19, 23–26). Interestingly, liver plays a central role in these adaptations. Hepatocytes promote tolerance by producing glucose (19, 21–23), degrading heme (18, 23), and responding to glucocorticoid (24) and GDF15 (25) as well as by secreting acute phase proteins (14, 27), FGF21 (26), and very low density lipoprotein (15, 25, 26). When managed well, the net effect is reduced organ damage, preserved heart function, and prevention of hypothermia (13, 15, 18, 19, 23, 25, 26, 28). Moreover, hepatocytes and other liver cells coordinate to promote disease resistance by destroying an invading pathogen, altering inflammation, and regulating lipoproteins that can eliminate/clear pathogens (14, 16, 17, 20, 27, 29–34). These defenses are important to avert life-threatening pathology, as patients with liver cirrhosis are more prone to acquire sepsis and less able to survive sepsis (27, 35, 36). Hence, liver may be a critical target organ to stimulate host defenses capable of improving sepsis outcomes.

Stress defense networks mediate damage control and promote adaptations that underlie disease tolerance (7, 9–12). Which networks in hepatocytes contribute to liver defenses against sepsis is unclear, but transcriptional programming is important for the process (37). Previous works identified a role for the transcription factors glucocorticoid receptor, farnesoid X receptor, and peroxisome proliferator-activated receptor alpha (24, 38–40). In this study, we considered whether stress defending transcription factors nuclear factor erythroid 2 related factor-1 (Nrf1) and -2 (Nrf2) contribute to such programming. Nrf1 and Nrf2 can regulate genes that protect against ROS, proteotoxicity, impaired cell metabolism, and organelle dysfunction (41–46). While a role for Nrf1 in sepsis has not been studied, whole body Nrf2 deficient mice were found to have greater mortality (47). With respect to liver defenses, we have shown combined deletion of hepatic Nrf1 and Nrf2, by itself, results in predisposition to mortality and transcript profiles that resemble exposure to the sepsis-inducing endotoxin lipopolysaccharide (LPS) (48) as well as reduced glucose (49) and high density lipoprotein (50) output. Based on the role of liver and its adaptive role in disease tolerance, we reasoned these effects may influence sepsis outcomes. To investigate, here we modulate hepatic Nrf1 and Nrf2 level in mice and inject them with LPS or live *Escherichia coli* (*E. coli*) to determine whether the actions of hepatic Nrf1 and Nrf2 play a role in liver defenses against sepsis.

## Results

### Reduced hepatic Nrf1, but not Nrf2, activity impairs survival in LPS-induced sepsis

Nrf1 and Nrf2 can regulate genes that protect against ROS and other types of stress that emerge in liver during sepsis (41, 42, 44, 45, 47, 48, 51–57). To determine the effect of sepsis on hepatic Nrf1 and Nrf2 activity, we examined liver and survival of mice injected with LPS from *E*. *coli* O111:B4, which has been extensively used for investigating sepsis (15, 22, 25, 26, 28, 38, 47). During a 72- hour survival study, 1 mg/kg LPS resulted in 100 % survival, 10 mg/kg LPS resulted in 64 % survival, and 20 mg/kg LPS resulted in 10 % survival (Figure 1A). These sub-lethal, low, and high LPS doses were used in further experiments. Using an established loss-of-function model (48–50), we verified the detection of Nrf1 and Nrf2 in liver nuclei (Supplemental Figure 1A) and found that LPS caused expected changes in liver, such as reduced growth hormone receptor (58, 59) 8-hours post-injection with 20 mg/kg LPS (Supplemental Figure 1B) and increased mRNA of acute phase proteins serum amyloid A1, *Saa1*, and haptoglobin, *Hp* (14), 8- and 24-hours post-injection with 1 mg/kg LPS (Supplemental Figure 1C). Next, we examined surrogate factors for hepatic Nrf1 and Nrf2 activity. Compared to PBS, 8-hours post-injection with 20 mg/kg LPS reduced Nrf1 and Nrf2 in liver nuclei, while increasing NFκB (Figures 1B and 1C). This corresponded with reduced liver expression of the genes encoding Nrf1 (*Nfe2l1*) and Nrf2 (*Nfe2l2*) (Supplemental Figure 1D) and target genes of both (48), *Gclc*, *Gclm*, *Gsta3*, and *Gstm1* (Figure 1D). 8- and 24-hours post-injection, the sub-lethal 1 mg/kg dose of LPS had little effect on liver mRNA for *Nfe2l1* and a trending reduction in *Nfe2l2* mRNA (Supplemental Figure 1E). But, re-analyzed RNA sequencing (RNA-seq) differential expression data comparing liver from mice 12-hours post-injection with 1 mg/kg LPS to untreated control, see Ganeshan et al. (15) and Supplemental Table 1, still support that LPS reduces expression of *Gclc*, *Gclm*, *Gsta3*, and *Gstm1* (Supplemental Figure 1F). Altogether, these findings indicate that hepatic Nrf1 and Nrf2 activity is acutely impaired in LPS-induced sepsis, particularly during exposure to a highly lethal dose.

**Figure I.**
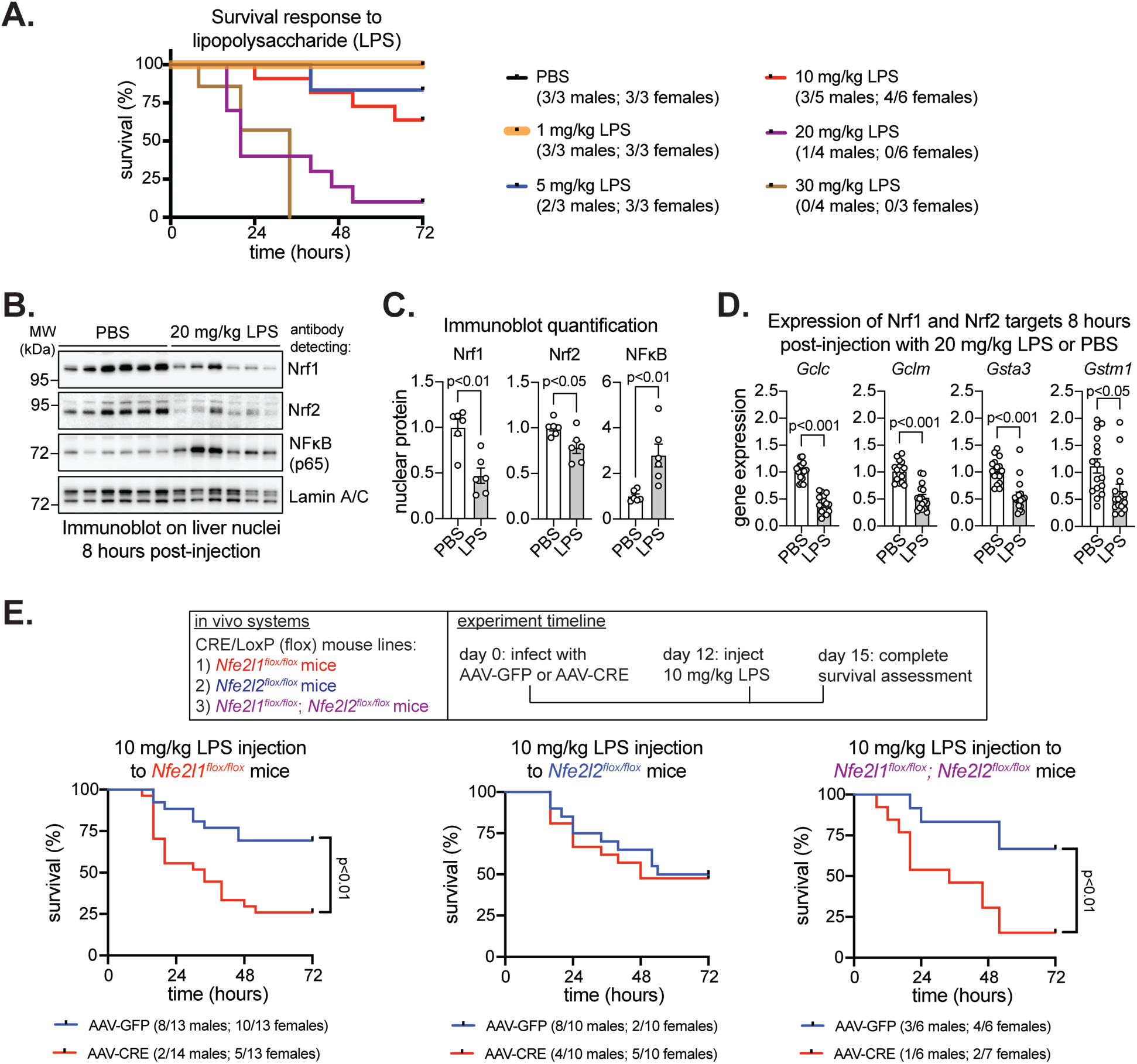
Hepatic Nrfl is important for surviving LPS-induced sepsis. A) Survival plot of mice injected with indicated doses ofLPS (n = 3-5 males; 3-6 females). B-C) Immunoblot (B) and its analysis (C), normalized by Lamin A/C, ofliver nuclei isolated 8-hours post injection with 20 mg/kg LPS (n = 3 males; 3 females). D) Expression ofNrfl and Nrf2 target genes in liver 8-hours post injection with 20 mg/kg LPS, normalized by 36b4 (n = 8 males; 8 females). E) Study design and survival plots of indicated tlox mice infected as indicated in panel (n = 6-14 males; 6-13 females). p-val ue in C and D determined by t-test. p-value in E determined by Log rank (Mantel-Cox) test. In A and E, number of survivors in each group is shown in symbol legend. Data in C and Dare mean± standard error of the mean, with individual data points shown.

The impact of ablating hepatic Nrf1 and Nrf2 activity days prior to sepsis onset on survival outcome was examined by utilizing mice with flox alleles for *Nfe2l1*, *Nfe2l2*, or both, which we described previously (48, 53). As before (48–50), *Nfe2l1^flox/flox^* mice, *Nfe2l2^flox/flox^* mice, and *Nfe2l1^flox/flox^*; *Nfe2l2^flox/flox^* mice were infected with adeno-associated virus (AAV) expressing Cre recombinase via thyroxine binding globulin promoter (AAV-CRE) to generate mice with hepatic deletion of Nrf1, Nrf2, or both, respectively. Controls received AAV expressing green fluorescent protein (AAV-GFP). As shown (Figure 1E), mice received AAV infection 12 days prior to LPS injection, consistent with previous (48–50). On day 12 (corresponding to ∼5 days of gene deficiency), mice were injected with 10 mg/kg LPS, a dose found to result in 64 % survival in regular mice (see Figure 1A). Survival was similar in controls, ranging from 50 % - 69 % between flox lines, and mice with hepatic Nrf2 deficiency were similar to control (survival: AAV-GFP = 50 % vs AAV-CRE = 45 %). In contrast, survival was reduced in mice with Nrf1 deficiency (survival: AAV-GFP = 69 % vs AAV-CRE = 26 %) and combined deficiency (survival: AAV-GFP = 58 % vs AAV-CRE = 23 %). Hence, we demonstrate that hepatic Nrf1, but not Nrf2, is required to protect against sepsis, and impairment to its activity, such as occurs in high LPS exposure, may predispose for lethality.

### Effects of hepatic Nrf1, Nrf2, and combined deficiency in LPS-induced sepsis

We next sought to identify inflammatory and metabolic factors that may explain reduced survival in hepatic Nrf1 deficient mice exposed to LPS. Previous works show hepatic Nrf1 can counteract liver inflammation, promote proteostasis, and regulate lipid and glucose metabolism (45, 46, 48–54). Here, compared to PBS, 10 mg/kg LPS robustly increased plasma levels of several inflammatory cytokines that underlie sepsis 8-hours post-injection, but hepatic deficiency for Nrf1, Nrf2, or both had no effect on these levels (Figure 2A and Supplemental Figure 2A) or on LPS-induced changes to plasma glucose (Figure 2B) and bile acids (Supplemental Figure 2B). In contrast, liver expression of the chemokine *Ccl2* was increased and the transcription factor *Pparα* was decreased by hepatic Nrf1 and combined deficiency (Supplemental Figure 2C), irrespective of LPS. Likewise, liver inflammation was increased in hepatic Nrf1 and combined deficiency, irrespective of LPS (Supplemental Figures 2D and 2E). Hepatic Nrf1 deficiency increased the unfolded protein response marker, phosphorylated eukaryotic initiation factor 2α (p-eIF2α), but had no effect on the protein translation activity marker, phosphorylated ribosomal protein S6 kinase (p-S6K), (Supplemental Figures 2F and 2G), suggesting these livers were under endoplasmic reticulum (ER) stress. Compared to PBS, LPS increased circulating triglyceride in control mice and in mice with hepatic Nrf2 deficiency, but this effect was blunted in the mice with hepatic Nrf1 deficiency and in those with combined deficiency (Figure 2C). Altogether, we show that reduced survival in hepatic Nrf1 deficient mice is not due to altered systemic inflammation but does correspond with increased ER stress and inflammation in the liver as well as lower levels of circulating triglyceride.

**Figure 2.**
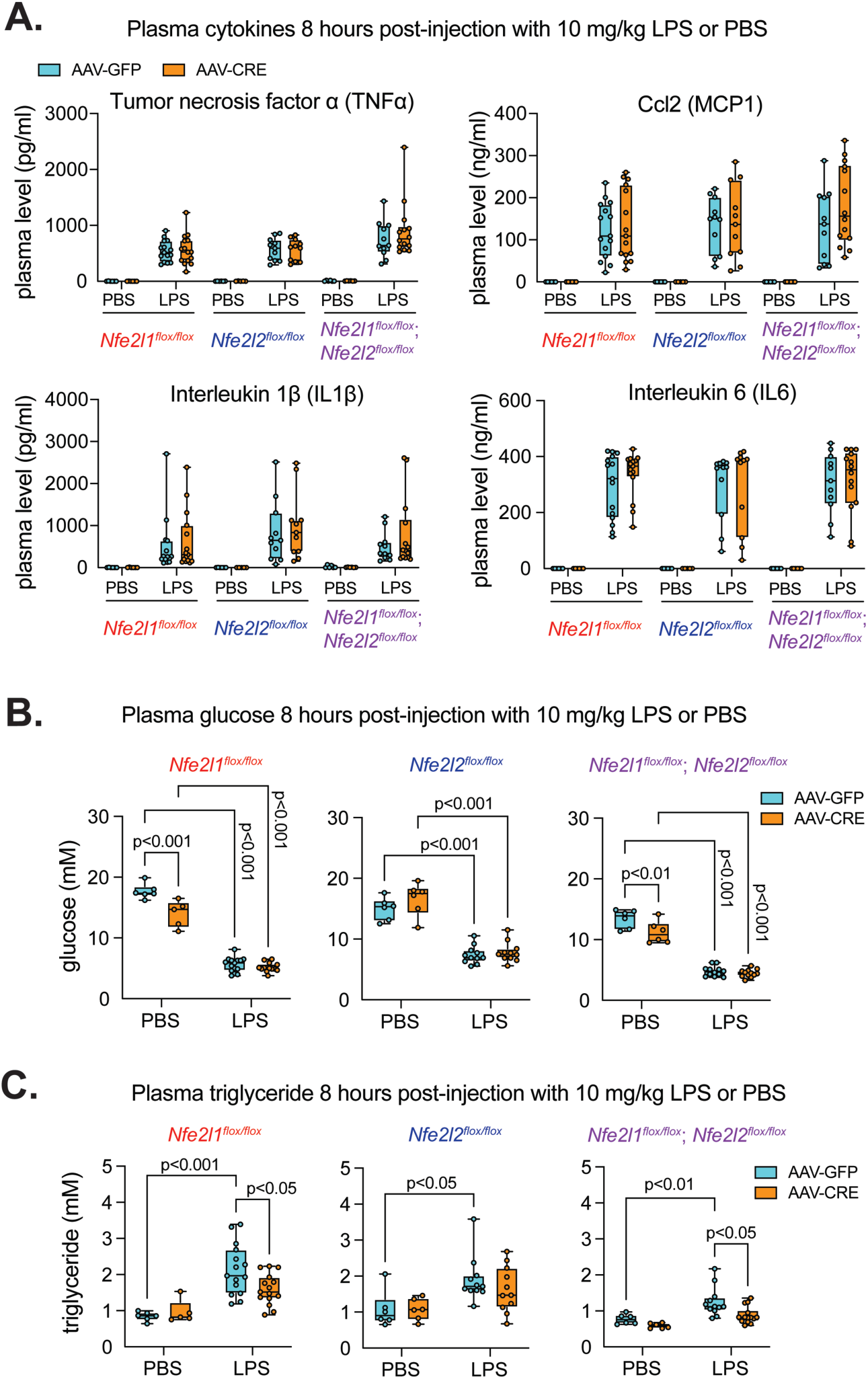
Hepatic Nrfl deficiency alters plasma triglyceride, but not cytokines or glycemia. A-C) Plasma cytokines (A), glucose (B), and triglyceride (C) for indicated tlox mice infected as indicated in panel (n = 2-7 males; 2-8 females). p-value in A determined by unpaired t-test, with adjustment for multiple comparisons. p-value in B and C determined by two-way ANOVA, with Tukey’s post-test. Data in A-Care box and whisker plots, with individual data points shown.

### Enhancing hepatic Nrf1 activity increases survival and triglyceride in LPS-induced sepsis

In complement to gene ablation (loss-of-function), we also investigated the effect of increasing hepatic Nrf1 activity (gain-of-function) prior to sepsis onset on survival outcome by comparing LPS responses of control mice infected with AAV-GFP to mice infected with an AAV expressing high levels of human NRF1 (AAV-hNRF1), which we described previously (48). In this case, we used 20 mg/kg LPS, a dose we previously found to result in only 10 % survival in regular mice (see Figure 1A). As shown (Figure 3A), 12 days after AAV infection (corresponding to ∼5 days of increased Nrf1 activity), mice were injected with LPS and a 72-hour survival study done. Compared to control, mice with hepatic Nrf1 gain-of-function had improved survival (survival: AAV-GFP = 11 % vs AAV-hNRF1 = 46 %). Hence, enhancing hepatic Nrf1 activity is protective against LPS-induced sepsis, which is the opposite effect of hepatic Nrf1 deficiency. Therefore, we show that hepatic Nrf1 is a critical factor involved in promoting liver defenses against sepsis.

**Figure 3.**
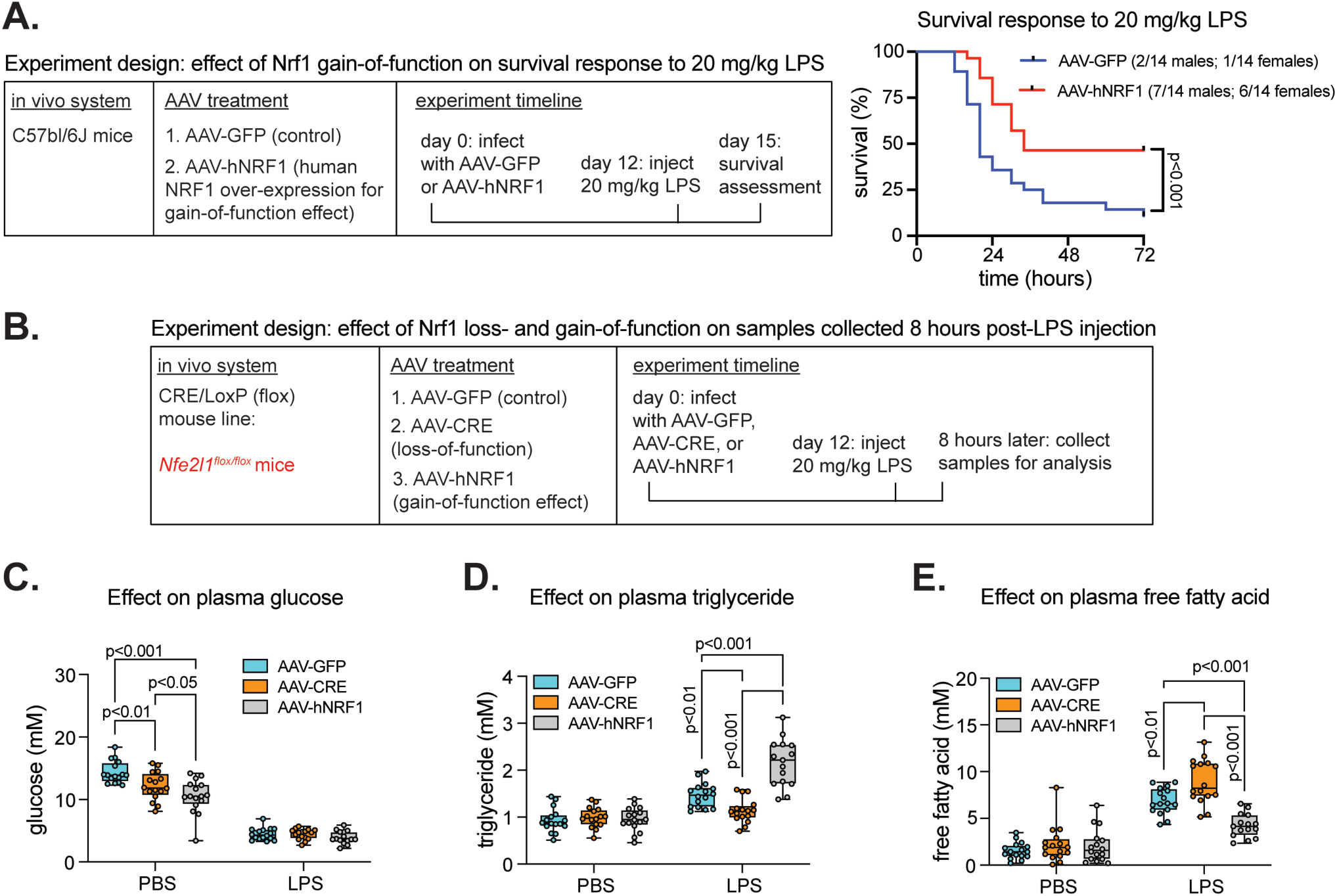
Increasing hepatic Nrfl activity improves survival during sepsis and affects plasma triglyceride and free fatty acid in an inverse manner to hepatic Nrfl deficiency. A) Study design and survival plot of mice infected withAAV-GFP or AAV expressing human NRF1 in hepatocytes (AAV-hNRF 1) and injected with 20 mg/kg LPS (n = 14 males; 14 females). Number of survivors in each group is shown in symbol legend. B) Study design comparing control to Nfe2llflox/flox mice with hepatic Nrfl loss- or gain-of-function 8-hours post injection with PBS or 20 mg/kg LPS. C-E) Corresponds to study design in (B). Plasma glucose (C), triglyceride (D), and free fatty acid (E) in PBS and LPS injected Nfe211flox/flox mice (n = 7-8 males; 8 females). p-value in A determined by Log rank (Mantel-Cox) test. p-value in C-E determined by two-way ANOVA, with Tukey’s post-test. Data in C-E are box and whisker plots, with individual data points shown.

Next, using *Nfe2l1^flox/flox^* mice, we used complementary AAV-induced models for hepatic Nrf1 loss-(AAV-CRE) and gain-of-function (AAV-hNRF1) to identify sepsis-related parameters affected in the same opposing pattern as survival outcomes, compared to control mice (AAV-GFP) 8 hours post-injection with 20 mg/kg LPS or PBS (schematic in Figure 3B). This was not the case for plasma glucose and ketones (Figure 3C and Supplemental Figure 3A). Moreover, hepatic Nrf1 loss-of-function decreased and gain-of-function increased plasma IGF1 in PBS injected but not in LPS injected mice (Supplemental Figure 3B) and the same pattern occurred for liver expression of growth hormone receptor in PBS and LPS injected mice (Supplemental Figure 3C). Liver expression of *Ccl2* was increased by hepatic Nrf1 loss-of-function, but unaffected by hepatic Nrf1 gain-of-function (Supplemental Figure 3D). Interestingly, in only the LPS injected mice, hepatic Nrf1 loss-of-function decreased and gain-of-function increased plasma triglyceride (Figure 3D), whereas hepatic Nrf1 loss-of-function increased and gain-of-function decreased plasma free fatty acid (Figure 3E). This shows the protective effect of hepatic Nrf1 activity against LPS-induced sepsis coincide with reduced free fatty acid and increased triglyceride in the bloodstream.

### Enhancing hepatic Nrf1 activity increases stress defense programming in LPS-induced sepsis

Given current understanding (42, 45, 46, 48, 50–54, 60, 61), we reasoned that hepatic Nrf1 activity protects against LPS-induced sepsis via gene regulation. To investigate, RNA-seq was done on livers from control (AAV-GFP), hepatic Nrf1 loss-of-function (AAV-CRE), and hepatic Nrf1 gain-of-function (AAV-hNRF1) *Nfe2l1^flox/flox^* mice 8 hours post-injection with 20 mg/kg LPS. Principle component analysis (PCA) comparing AAV-CRE to AAV-GFP and AAV-hNRF1 to AAV-GFP revealed hepatic Nrf1 loss- and gain-of-function did not cause major changes to transcript profiles (Supplemental Figures 4A and 4B). This was not surprising, given that LPS on its own drives broad and robust alterations to liver transcript profiles (15, 37, 62), and this may mask effects of altering hepatic Nrf1 activity. Interestingly, in previous work on liver in less stressful conditions (48, 51–53) hepatic Nrf1 loss-of-function has been shown to alter expression of thousands of genes, but there were no genes affected by hepatic Nrf1 loss-of-function in this case (Supplemental Figure 4A) when liver is experiencing stress caused by LPS exposure and which, by itself, can impair hepatic Nrf1 activity (see Figures 1B-1D). In contrast, compared to control using false discovery rate p-value adjusted DESeq2 analysis, expression of 330 genes were significantly different (69 down-regulated and 261 up-regulated) in liver of mice with hepatic Nrf1 gain-of-function (Supplemental Figure 4B and Supplemental Tables 2 and 3). Hence, hepatic Nrf1 gain-of-function impacted hepatic transcriptional programming, and this may underlie protection against sepsis.

Ingenuity pathway analysis (IPA) was done to identify pathways most affected by hepatic Nrf1 gain-of-function. Genes undergoing IPA were selected using Wald statistics ≥ 3 (391 genes) and ≤ -3 (76 genes), which amounted to 467 genes. Focusing on diseases-and-biofunctions, pathways most affected include “necrosis”, “oxidative stress”, “inflammation of body cavity”, and “organismal death”, and molecular activity prediction analysis suggest these pathways were less active in liver with hepatic Nrf1 gain-of-function (Figure 4A). A heatmap of transformed RNA-seq values for genes corresponding to the “necrosis” pathway was also done (Figure 4B), and show increased mRNA level of several genes that counteract cell stress and many of which are known to be regulated by Nrf1, such as *Gclm*, *Gstm1*, *Psma4*, *Psmb5*, and *Vcp* (42, 45, 48, 51, 52, 54, 60).

**Figure 4.**
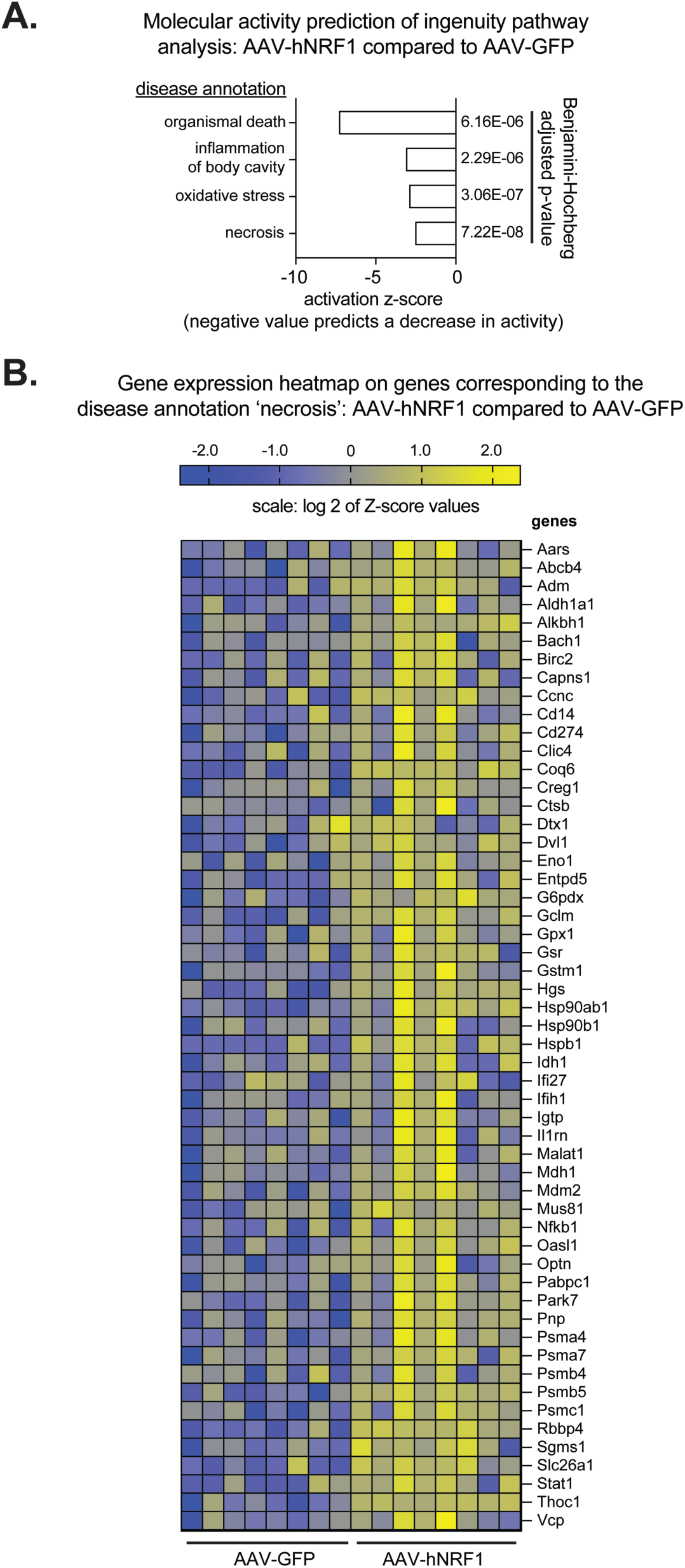
Effect of increasing hepatic Nrfl activity on liver gene expression during sepsis. A) Compared to control mice (AAV-GFP), pathways enriched in liver ofNfe21lflox/flox mice with hepatic Nrfl gain-of-function (AAV-hNRFl) 8-hours post injection with 20 mg/kg LPS, (n = 4 males; 4 females). p-value determined by Benjamin Hochberg analysis. B) Heatmap of relative expression for genes corresponding to pathway disease annotation: necrosis. Shown are log 2 values after transforming to Z-score.

Altogether, this suggests enhancing hepatic Nrf1 activity increased hepatic resistance to sepsis-induced stress, and this may directly or indirectly promote liver defenses that protect against sepsis.

### Hepatic Nrf1 activity protects against *E. coli*-induced sepsis and hypothermia

To ensure results using LPS translate to bacterial inflammation, we investigated whether hepatic Nrf1 loss- and gain-of-function had the same opposing effects on survival and sepsis-related parameters in *Nfe2l1^flox/flox^*mice infected with live *E. coli*. An *E. coli* dose response was done (Supplemental Figure 5A), from which we determined that administering 4.0 x 10^8^ colony forming units (CFU) *E. coli* in males and 3.2 x 10^8^ CFU *E. coli* in females results in approximately 50 % survival during a 120-hour survival study. Compared to PBS, *E. coli*-induced sepsis reduced liver expression of Nrf1 and Nrf2 target genes 24-hours post-infection (Supplemental Figure 5B), similar to the effect of high dose LPS (Figures 1B-1D). In comparison to control, hepatic Nrf1 loss-of-function reduced survival and hepatic Nrf1 gain-of-function increased survival (Figure 5A). Similar to the LPS model (Figures 2B and 3C), altering hepatic Nrf1 activity had no effect on sepsis-induced changes to blood glucose (Figure 5B) whereas plasma triglyceride was increased in mice with hepatic Nrf1 gain-of-function (Figure 5C) 24 hours post-infection. We also monitored body temperature prior to (i.e., time 0) and 24 hours post-infection, as hypothermia is a critical trigger for sepsis-induced lethality and the liver is capable of defending against this trigger by fueling thermogenic tissues with VLDL-triglyceride (15, 25, 26, 63). As with the survival outcomes (Figure 5A), hepatic Nrf1 loss-of-function worsened hypothermia and gain-of-function improved hypothermia, compared to control mice, whereas there was no body temperature difference prior to infection (Figure 5D). Thus, we show hepatic Nrf1 activity protects against *E. coli*-induced sepsis, with corresponding changes in circulating triglyceride, and also that the mechanism of this protection may involve liver defense against hypothermia.

**Figure 5.**
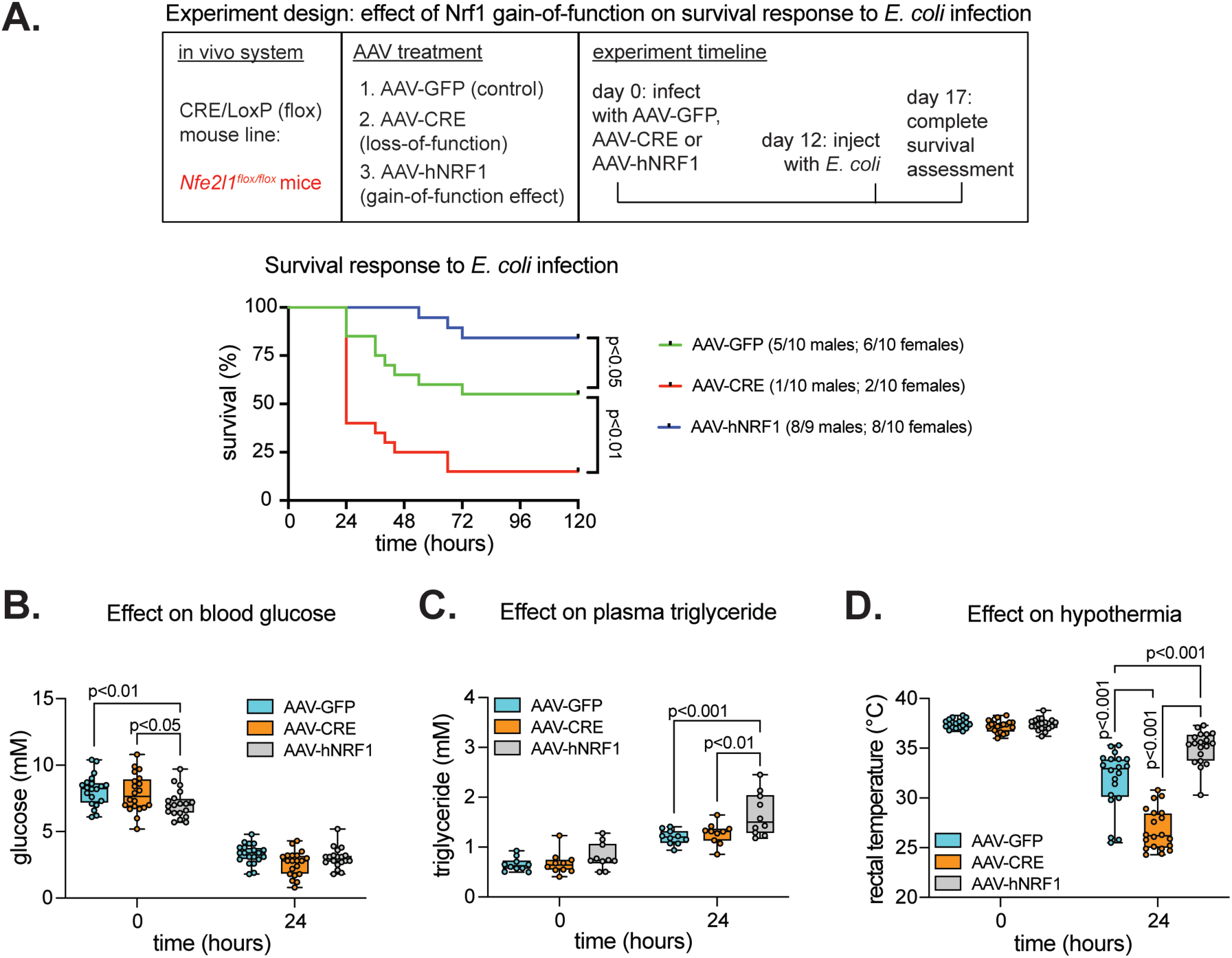
Hepatic Nrfl is important for surviving E. coli-induced sepsis, corresponding with increasing plasma triglyceride and protection against hypothermia. A) Study design and survival plot comparing control (AAV-GFP) to Nfe2llflox/flox mice with hepatic Nrfl loss-(AAV-CRE) or gain-of-function (AAV-hNRFl) in response to 4.0 x 10_8_ CFU E.coli (n = 9-10 males) or 3.2 x 10_8_ CFU E.coli (n = 10 females). Number of survivors in each group is shown in symbol legend. B-D) Comparison of control to Nfe21lflox/flox mice with hepatic Nrfl loss- or gain-of-function 24-hours post-injection ofE. coli. Shown is glucose (B) in blood (n = 10 males; 9-10 females), triglyceride (C) in plasma (n = 5 males; 5 females), and rectal temperatures (D) assessing hypothermia **(n** = 10 males; 9-10 females). p-value in A determined by Log rank (Mantel-Cox) test. p-value in B-D determined by two-way ANOVA with Tuk:ey’s post-test. Data in B-D are box and whisker plots, with individual data points shown.

### Hepatic Nrf1 activity protects against sepsis and hypothermia by promoting VLDL secretion

Triglyceride rich VLDL can promote survival in mice with sepsis by counteracting hypothermia (15, 25, 26). So, we investigated whether reciprocal changes in circulating triglyceride that occur in hepatic Nrf1 loss- and gain-of-function mice (Figures 2, 3, and 5) is causally linked to coinciding effects on liver defense against sepsis-induced hypothermia. First, we injected the sub-lethal dose of 1 mg/kg LPS (see Figure 1A) to *Nfe2l1^flox/flox^* mice infected 12 days prior with AAV-GFP (control), AAV-CRE (loss-of-function), or AAV-hNRF1 (gain-of-function) and monitored body temperature, sepsis score, plasma triglyceride and other sepsis-related parameters 0-, 8-, and 24- hours post-injection (see Figure 6A). There were changes to some plasma cytokines (Supplemental Figure 6A), liver inflammation (Supplemental Figures 6B and 6C), and plasma glucose (Supplemental Figure 6D), but this did not follow a consistent pattern that can explain the outcomes. Similarly, hepatic Nrf1 gain-of-function increased liver expression of *Ghr* 0-, 8-, and 24-hours post-injection (Supplemental Figure 6E) but hepatic Nrf1 loss-of-function had no effect, whereas hepatic Nrf1 loss-of-function reduced plasma IGF1 0- and 24-hours post-injection (Supplemental Figure 6F), but hepatic Nrf1 gain-of-function had no effect. In contrast to these inconsistent patterns, hepatic Nrf1 loss-of-function worsened whereas gain-of-function improved hypothermia and the sepsis score (Figures 6B and 6C), which coincided with reduced plasma triglyceride in hepatic Nrf1 loss-of-function mice and increased plasma triglyceride in hepatic Nrf1 gain-of-function mice (Figure 6D). Hence, we confirm there is a corresponding relationship between sepsis severity, body temperature, and circulating triglyceride with hepatic Nrf1 activity. Liver-derived circulating triglyceride primarily originates from secreted VLDL, and this lipoprotein is capable of mediating disease tolerance (15, 25, 26, 28). So, we investigated whether hepatic Nrf1 promotes VLDL secretion to protect against sepsis severity and hypothermia in LPS exposed mice. First, we performed a VLDL secretion assay. 16-hours post-injection with 1 mg/kg LPS, mice were co-injected with lipoprotein lipase inhibitor poloxamer 407 and then triglyceride was monitored 0-, 2-, and 4-hours later. Compared to control, by hour 4 plasma triglyceride was reduced in mice with hepatic Nrf1 loss-of-function and increased in mice with hepatic Nrf1 gain-of-function (Figure 6E), showing hepatic Nrf1 activity positively correlates with VLDL secretion. Second, we investigated whether increased VLDL secretion in hepatic Nrf1 gain-of-function mice is causally linked to protection against sepsis by employing use of the VLDL secretion inhibitor lomitapide, which is a clinically used agent that inhibits triglyceride incorporation into VLDL (64). As expected, lomitapide blunted plasma triglyceride levels in AAV-GFP and AAV-hNRF1 infected mice (Supplemental Figure 6G). Control (AAV-GFP) and hepatic Nrf1 gain-of-function (AAV-hNRF1) mice were injected with LPS + vehicle or LPS + lomitapide, whereas the hepatic Nrf1 loss-of-function (AAV-CRE) mice were injected with only LPS + vehicle. Body temperature and sepsis score was monitored 0-, 8-, and 24-hours post-injection (Figures 6F and 6G). As before, hepatic Nrf1 loss-of-function mice had more severe hypothermia and sepsis score than LPS + vehicle treated control mice. Conversely, hepatic Nrf1 gain-of-function mice treated with LPS + vehicle had improved hypothermia and sepsis score than LPS + vehicle treated control mice. However, the beneficial effect of hepatic Nrf1 gain-of-function was lost in mice treated with LPS + lomitapide, and this was so severe that body temperature and sepsis score in either control or hepatic Nrf1 gain-of-function mice treated with LPS + lomitapide were now similar to that found in hepatic Nrf1 loss-of-function mice treated with LPS + vehicle. Hence, we show that enhancing hepatic Nrf1 activity causes an increase in the secretion of triglyceride rich VLDL during sepsis and that this effect is required for hepatic Nrf1 activity to protect against sepsis pathology.

**Figure 6.**
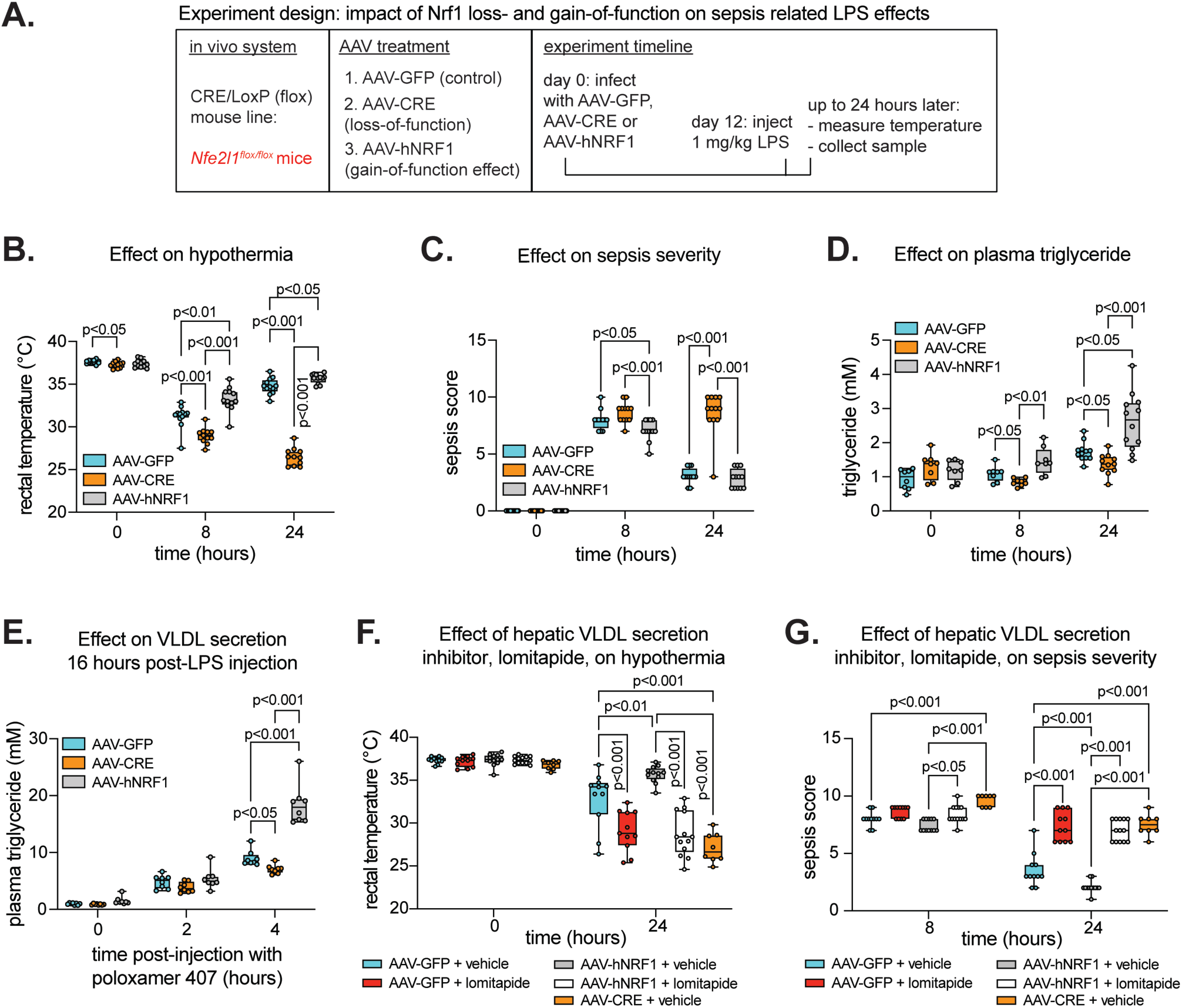
Hepatic Nrfl promotes VLDL-dependent liver defenses that protect against sepsis severity and hypothermia. A) Study design comparing control (AAV-GFP) to Nfe211flox/flox mice with hepatic Nrfl loss­(AAV-CRE) or gain-of-function (AAV-hNRFl) 0-24 hours post-injection with **1** mg/kg LPS. B-D) Rectal temperature (B), sepsis score (C), and plasma triglyceride (D) 0-24 hours post-injection with LPS (n = 4-6 males; 4-6 females). E) VLDL secretion 16 hours post-injection with LPS. Plasma triglyceride was measured 0-4 hours after injecting 1 g/kg p-407, which prevents clearance of circulating triglyceride by inhibiting lipoprotein lipase (n = 4 males; **4** females). F-G) Rectal temperature **(F)** and sepsis score (G) 0-24 hours post-injection with LPS ± co-injection with 2 mg/kg of hepatic VLDL secretion inhibitor, lomitapide (n = 4-6 males; 4-7 females). p-value in B-G determined by two-way ANOVA with Tukey’s post-test. Data in B-G are box and whisker plots, with individual data points shown.

## Discussion

Sepsis is a major health challenge and there are limited interventional options (1–6). One pressing need is a strategy for reducing tissue damage to preserve organ function and improve recovery (6). Recent works show there are endogenous processes capable of this feat, referred to as disease tolerance (7–10). There is interest in stimulating mechanisms of disease tolerance to improve sepsis outcomes (2, 6–10). Mechanisms underlying disease tolerance are incompletely understood, but mediators of tissue damage control and their effect on systemic metabolism are considered critical components (7, 9–11). Here, we investigated whether transcription factors Nrf1 and Nrf2 in hepatocytes, which regulate genes involved in stress defense (41–46), protect against sepsis. We demonstrate that ablating hepatic Nrf1, but not Nrf2, predisposes for mortality whereas genetic-induction of hepatic Nrf1 activity improves survival in mice with LPS- and *E. coli*-induced sepsis. Thus, we identify hepatic Nrf1 as an important factor for liver defense against sepsis.

The liver plays a central role in defending against sepsis and hepatocytes contribute to this role, which includes adaptations in glucose, lipid, bile acid, ketone, and lipoprotein metabolism as well as influences on inflammation (14–36, 38–40). We sought to determine whether effects of altered hepatic Nrf1 activity on survival in sepsis was linked to these parameters. Compared to control, hepatic Nrf1 loss- and gain-of-function did not alter systemic inflammation in mice with severe sepsis and had no effect on sepsis-induced reductions in glycemia. Also, complementary alterations to hepatic Nrf1 activity had parallel, not opposing, effects on ketones and no effect on bile acids in *Nfe2l1^flox/flox^* mice. Liver inflammation was increased in hepatic Nrf1 loss-of-function mice, and this was elevated 24-hours post-injection with sub-lethal sepsis, but the opposite did not occur in hepatic Nrf1 gain-of-function mice. Thus, the effect of altered hepatic Nrf1 activity on survival was not linked to the effects of sepsis on glucose, ketone, and bile acid metabolism as well as systemic inflammation. Since glucocorticoid receptor, farnesoid X receptor, and peroxisome proliferator-activated receptor can protect against sepsis by regulating these metabolic processes (24, 38–40), hepatic Nrf1 may control an independent program that operates in parallel with these other factors. We did find a positive but inconsistent correlation for hepatic Nrf1 activity with liver growth hormone receptor and circulating IGF1, indicating there may be a link with the growth hormone/IGF1 axis. We previously found this link in a non-sepsis context (49) but are uncertain of the significance. In contrast, we consistently identified a positive correlation for hepatic Nrf1 activity with circulating triglyceride in sepsis, that hepatic Nrf1 loss-of-function mice have reduced whereas hepatic Nrf1 gain-of-function mice have increased VLDL secretion, and that inhibiting VLDL secretion blocked protective effects of enhancing hepatic Nrf1 activity against hypothermia and sepsis severity. Hence, our results show that hepatic Nrf1 protects against sepsis, at least in part, by promoting an increase in the secretion of triglyceride rich VLDL.

VLDL protects against sepsis by fueling thermogenesis to avert hypothermia (15, 25, 26) and by facilitating pathogen elimination (30–32, 34). We show that Nrf1-regulated VLDL secretion protects against hypothermia-linked mortality (63). While we did not delineate the mechanism by which hepatic Nrf1 controls VLDL secretion, we did examine the effect of hepatic Nrf1 on transcript profiles. RNA-seq results show hepatic Nrf1 gain-of-function liver has increased stress defense gene expression, consistent with previous (42, 46, 48, 50–54), which may reduce liver susceptibility to damage. Indeed, hepatic Nrf1 loss-of-function liver had increased ER stress and inflammation. So, while we cannot draw a direct link, we speculate that liver with enhanced hepatic Nrf1 activity is better suited to manage the stress of sepsis and, in turn, to mediate defenses that promote survival. Consistent with this, we found that severe sepsis reduced Nrf1 and Nrf2 in liver nuclei, coinciding with reduced target gene expression, indicating susceptibility to mortality in severe sepsis may, in part, result from impaired hepatic Nrf1 and Nrf2 activity. Indeed, we have shown that combined deletion of hepatic Nrf1 and Nrf2 in non-sepsis adult mice, by itself, results in a 36 % mortality rate just 28-days after inducing the gene deletion (48).

By identifying a protective role for hepatic Nrf1 in sepsis, we uncover four avenues for future research. First, we mainly use LPS here, due to its prior uses investigating sepsis and disease tolerance (15, 22, 25, 26, 28, 38, 47). Though we confirmed key results using *E. coli*-induced sepsis, further work is needed to determine whether altering hepatic Nrf1 activity has similar effects in other types (e.g., polymicrobial, gram positive bacteria, virus, and fungal) and modes (e.g., intravenous versus peritonitis) of infection to ascertain effectiveness in protecting against sepsis. Second, transcriptional programming is important for counteracting sepsis (24, 37–40). So, it is critical to delineate which genes Nrf1 regulates to protect against sepsis, as this may reveal strategies for stimulating disease tolerance. Third, it is important to determine the mechanism by which Nrf1 is impaired in sepsis and whether reversing this effect can improve sepsis outcomes. This may also be the case for Nrf2. Though hepatic Nrf2 was not required to protect against sepsis, Nrf2 in other cell types and its induction has been shown to be protective (47, 65, 66) and may require hepatic Nrf2 to do so. Fourth, further insight is needed regarding the impact that hepatic Nrf1 has on the quantity and quality of VLDL and other sepsis counteracting lipoproteins such as high-density lipoprotein (17, 31, 32, 50). Likewise, further research on the mechanism by which lipoproteins protect against sepsis is likely to reveal important insight to treat sepsis.

In summary, we show hepatic Nrf1 protects against sepsis via promoting VLDL secretion, which can prevent hypothermia to reduce sepsis severity and promote survival. Our finding underscores the value of investigating stress defense programming in liver and its impact on the mechanisms underlying tissue damage control in sepsis.

## Methods

### Animal used in study

Experiments were done on 20–30-week-old male and female mice on the C57BL/6J background. Mice were group housed at 21 °C on a 12 h light/dark cycle and provided *ad libitum* access to chow from LabDiet (Prolab RMH 3000; catalog#: 5058) and water. Mice containing flox alleles in the genes for Nrf1 (*Nfe2l1^flox/flox^*), Nrf2 (*Nfe2l2^flox/flox^*), or both Nrf1 and Nrf2 (*Nfe2l1^flox/flox^*; *Nfe2l2^flox/flox^*) were bred in-house and described previously (48, 53).

To induce hepatocyte-specific loss-of-function for Nrf1 activity, Nrf2 activity, or both, recombination of respective flox alleles was induced by retro-orbital infection of mice, while under anesthesia (3% isoflurane at oxygen flow rate of 1 L/minute), with a serotype 8 adeno-associated virus (AAV) that expresses Cre recombinase via the hepatocyte-specific thyroxin binding globulin promoter (AAV-CRE). To induce hepatocyte-specific gain-of-function for Nrf1 activity, mice were infected with AAV expressing human NRF1 (AAV-hNRF1). In each case, littermate control mice were infected with AAV expressing green fluorescent protein (AAV-GFP). Use of these AAVs were established by us previously (48–50). In each experiment, males were infected with 3.0 x 10^11^ AAV particles and females with 2.0 x 10^11^ AAV particles, as this was found to result in more equivalent expression between sexes. All experiments involving AAVs were done within 17 days of infection. AAV-CRE (AAV8.TBG.PI.Cre.rBG) and AAV-GFP (AAV8.TBP.PI.eGFP.WPRE.bGH) were acquired from Addgene (#107787-AAV8 and #105535-AAV8). AAV-hNRF1 was produced by the University of Pennsylvania Vector Biocore and is described in our previous work (48).

### Sex as a biological variable

Male and female mice were used throughout. All between-sex results were in good agreement and so male and female samples were pooled when performing statistical analysis.

### Sepsis induction and assessment of severity

Sepsis was induced in mice via intraperitoneal (IP) injection with either lipopolysaccharide (LPS) from *E*. *coli* O111:B4 (Sigma, catalog #L2630) or with live DH5α *Escherichia coli* (*E*. *coli*; New England Biolabs, catalog #C2987H). Doses are described in the text, figures, and figure legends. In experiments that employed AAVs, sepsis was induced on day 12 after AAV administration. Mice were monitored for up to 72 hours when using LPS and 120 hours when using *E. coli*. Monitoring for symptoms and assignment of a murine sepsis score was done according to the guidelines by Shrum et al. (67). Body temperature was measured using a rectal probe thermometer (BIOSEB Lab Instruments, catalog #BIO-TK8851). Mice were euthanized upon reaching a predefined humane endpoint and, at that point, determined to have not survived the sepsis challenge.

### Bacteria preparation for sepsis induction

*E. coli* were heat shocked in a water bath at 42 °C for 30 seconds, placed on ice for 2 minutes, and grown to log phase in LB broth (Fisher, catalog #BP1426) at 37 °C in a shaking incubator. *E. coli* growth curve was quantified by correlating optical density at 600 nm to colony forming units (CFU), which was determined by spreading on LB agar plates (Fisher, catalog #BP1423). Prior to infecting mice, *E. coli* underwent centrifugation at 2000 g for 10 minutes at 4 °C, washed once with calcium and magnesium free phosphate buffered saline (PBS), and then resuspended in PBS.

### VLDL secretion assay and use of VLDL secretion inhibitor

Mice underwent a very low-density lipoprotein (VLDL) secretion assay to examine the role of Nrf1 and hepatic triglyceride output in protecting against sepsis. For this, mice were injected with a sub-lethal 1 mg/kg dose of LPS, fasted overnight, and 16 hours later received IP injection with 1g/kg of lipoprotein lipase inhibiting agent, poloxamer 407 (Sigma, catalog #16758). Blood was collected immediately prior to and up to 4 hours after receiving poloxamer 407 to measure plasma levels of triglyceride, as a surrogate readout for VLDL output. In addition to VLDL secretion assay, assessment of the effect of inhibiting VLDL secretion on body temperature and murine sepsis score post-LPS exposure was also done. In this case, prior to IP injection with LPS, mice were fasted overnight and received IP injection on the contralateral side with 2 mg/kg of lomitapide (Sigma, catalog #SML1385) in 10% DMSO/PBS, which suppresses VLDL secretion by inhibiting triglyceride loading into newly synthesizing VLDL particles in hepatocytes (64).

### Tissue collection

Tissues were collected from mice anesthetized via 3 % isoflurane at oxygen flow rate of 1 L/minute and euthanized by exsanguination and cervical dislocation. Blood was collected using a 26-gauge syringe needle via cardiac puncture and placed into an ethylenediaminetetraacetic acid (EDTA) containing tube (5 mM final concentration). Plasma was separated via centrifugation at 8000 g at 4 °C for 10 minutes, placed into a fresh tube, and snap frozen on dry ice. Liver was collected by initially flushing the vasculature of euthanized mice with PBS. The liver was excised, and a piece was immediately processed for histological analysis, nuclear isolation, or snap frozen on dry ice, as described in text and figure legends. All tissues were stored at -80 °C until further use.

### Plasma analysis

Plasma triglyceride levels were determined using Infinity Triglyceride reagent (ThermoFisher Scientific, catalog #TR22421) and a Synergy HT microplate reader. Plasma glucose was determined using an Accu-Check glucometer (Roche). Plasma cytokine levels were measured by Eve Technologies (https://www.evetechnologies.com/). Plasma IGF1 levels were determined by ELISA according to manufacturer’s instructions (R&D Systems, catalog #DY791).

### Histological analysis

A piece of the left liver lobe was fixed in 10 % neutral buffered formalin for 24 hours, subjected to three 24-hour washes in 70 % ethanol, embedded in wax, sectioned at 5 μm thickness, mounted on glass slides, and stained with hematoxylin and eosin (H&E). Images were capture using Aperio Scanscope CS image analysis System and Aperio Imagescope viewing software (version 12).

### Gene expression analysis using qPCR

Ribonucleic acid (RNA) were extracted from frozen liver tissue homogenized in TRIzol Reagent (ThermoFisher Scientific, catalog #15596018). RNA clean-up was performed using a Qiagen RNeasy plus column (ThermoFisher Scientific, catalog #74034). RNA concentration and quality were determined using a NanoDrop One spectrophotometer (Thermo-Fisher Scientific). One μg of RNA were reverse transcribed into complementary deoxyribonucleic acid (cDNA) using a Maxima First strand cDNA synthesis kit, with dsDNase (ThermoFisher Scientific, catalog #K1672). Gene expression was determined using quantitative polymerase chain reaction (qPCR) on cDNA template with a Bio Rad CFX384-well Real-time PCR Detection System and PowerUp SYBR green Master Mix (ThermoFisher Scientific, catalog #A25742). A list of primer sequences for measuring gene expression are provided in Supplemental Table 4. Change in expression of target genes were normalized to the reference genes *36b4* and *Tbp* and all data presented were in good agreement. Gene expression data are presented as relative expression compared to the control group after normalization, as indicated in the figure legends.

### Gene expression analysis using RNA sequencing

RNA sequencing (RNA-seq) was performed by the University of Saskatchewan Next Generation Sequencing Facility. As in previous (48), 500 ng of RNA per sample were used to prepare sequencing libraries using TruSeq Stranded mRNA Library Prep Kit from Illumina (catalog #20020595 and 20022371). Libraries were quantified using the Qubit 4 fluorometer and dsDNA BR DNA assay (Invitrogen, catalog #Q32850). The fragment size of each library was determined using the DNA Screentape (D1000) assay on the TapeStation 4150 instrument. Libraries were pooled and sequenced using a NextSeq 500/550 High Output Kit v2.5 (75 cycles), which generated 38 bp paired end reads. Reads were extracted and adapter trimmed using Illumina fastp (68). Reads were aligned using STAR version 2.7.9a(69), using the Gencode M32 mouse genome as reference. Generate FASTQ BaseSpace pipeline (version 1.1.0.64). There were > 25 million aligned reads per sample. Principal component analysis (PCA) was done to visualize sample clustering. Differential gene expression was determined using DESeq2 (version 1.22.2) (70).

### Nuclear fraction isolation

Liver nuclei was isolated to assess Nrf1 and Nrf2 levels, as we did previously(48). Approximately 250 mg of fresh liver from two mice were pooled (500 mg total), chopped into 1-3 mm pieces using a scalpel blade, and then homogenized with a Teflon homogenizer in STM buffer (250 mM sucrose, 50 mM Tris (pH 7.4), 5 mM MgCl_2_) containing 1 mM phenylmethylsulfonyl fluoride and 1mM dithiothreitol. Samples underwent 800 g centrifugation for 15 minutes at 4 °C. The pellet was resuspended in 10 ml of ice-cold buffer A (10 mM HEPES (pH 7.9), 10 mM KCl, 1.5 mM MgCl2, 0.34 M sucrose, 10 % glycerol, 1 mM DTT, 0.1 % Triton X-100) and 1x protease inhibitor cocktail (Roche, catalog #11697498001) and incubated on ice for 10 minutes. Pellets were then collected and washed once with ice-cold buffer A. Samples were centrifuged at 4 °C at 800 g for 15 minutes, supernatants were removed, and nuclear lysates were prepared for protein analysis by immunoblot in 1 ml ice-cold RIPA buffer containing 1x protease inhibitor cocktail.

### Immunoblot

Proteins were extracted from frozen liver pieces or nuclear fractions using RIPA buffer with fresh protease inhibitor cocktail. Concentrations were determined using a Pierce BCA protein assay kit (ThermoFisher Scientific, catalog #23250) and then heated at 95 °C for 5 minutes in loading buffer containing 1 % sodium dodecyl sulfate and then cooling on ice. 10 μg of protein was loaded onto a Novex 4-12 % Bis-Tris gel (ThermoFisher Scientific, catalog #NP0336), separated by size via electrophoresis in NuPAGE MOPS Running Buffer (ThermoFisher Scientific, catalog #NP0336), and transferred to a nitrocellulose membrane (Bio-Rad, catalog #1620115). Membranes were blocked in tris-buffer saline containing 0.1 % tween 20 (TBST) and 5 % powdered skim milk, and then incubated overnight at 4 °C with primary antibodies targeting the protein of interest. Antibodies used are described in Supplemental Table 5. Membranes were washed with TBST and incubated with horse radish peroxidase-conjugated secondary antibodies at room temperature for 60 minutes. Membranes were washed with TBST and levels of immuno-detected protein measured by incubating in Super Signal West Femto Maximum Sensitivity Substrate (ThermoFisher Scientific, catalog #34096) to generate chemiluminescence, which was detected using a BioRad ChemiDoc MP Imaging System. Level of signal was analyzed using ImageJ (https://imagej.net/ij/).

### Quantification and statistical analysis

Statistical analysis of RNA-seq data was done using DESeq2 (70). All other data was analyzed using GraphPad PRISM (version 10.2.2). Significance was defined as p<0.05, using the Mantel-Cox Log rank test, t-test, one-way, or two-way analysis of variance (ANOVA) and Dunnett’s or Tukey’s post-test, as indicated in figure legends.

## Resource availability, author contributions, and funding acknowledgment

### Supplemental Information

Supplemental information is available for this manuscript. This includes the following: Supplemental Figures 1-6 and Supplemental Tables 1-5.

### Study approval

Animal studies were approved by the University of Saskatchewan’s Animal Care Committee under protocol number AUP20180090.

### Data Availability

Upon acceptance for publication, data corresponding to figures will be publicly available to view in the Mendeley repository and raw sequence reads pertaining to RNA-seq data will be publicly available to view in the National Center for Biotechnology Information (NCBI) repository. An excel file of data values is provided also.

### Author Contributions

Designing research studies, MJT and SBW; Conducting experiments and acquiring data, MJT, LL, MGA, AH, and SBW; analyzing data, MJT, LL, and SBW; writing - original draft, MJT and SBW; writing - review and editing, SBW.

### Declaration of Interests

The authors declare there are no competing interests.

### Funding Sources

Grants supporting this research were awarded to S.B.W. from the Canadian Institutes of Health Research (project #PJT-174988). M.J.T., M.G.A., and A.H. were supported by a scholarship from the University of Saskatchewan’s College of Medicine. A.H. was also supported by a scholarship from the Natural Sciences and Engineering Research Council of Canada. S.B.W. was supported by a New Investigator Award from the Heart and Stroke Foundation of Canada.

## Notes

### Competing Interest Statement

The authors have declared no competing interest.

### Summary of Updates

This manuscript has been revised to provide more details on the Methods, and more supplementary files.

