## Supplemental Figures S1-S6 for "Hepatic Nrf1 (*Nfe2l1*) promotes VLDL dependent liver defense against sepsis"

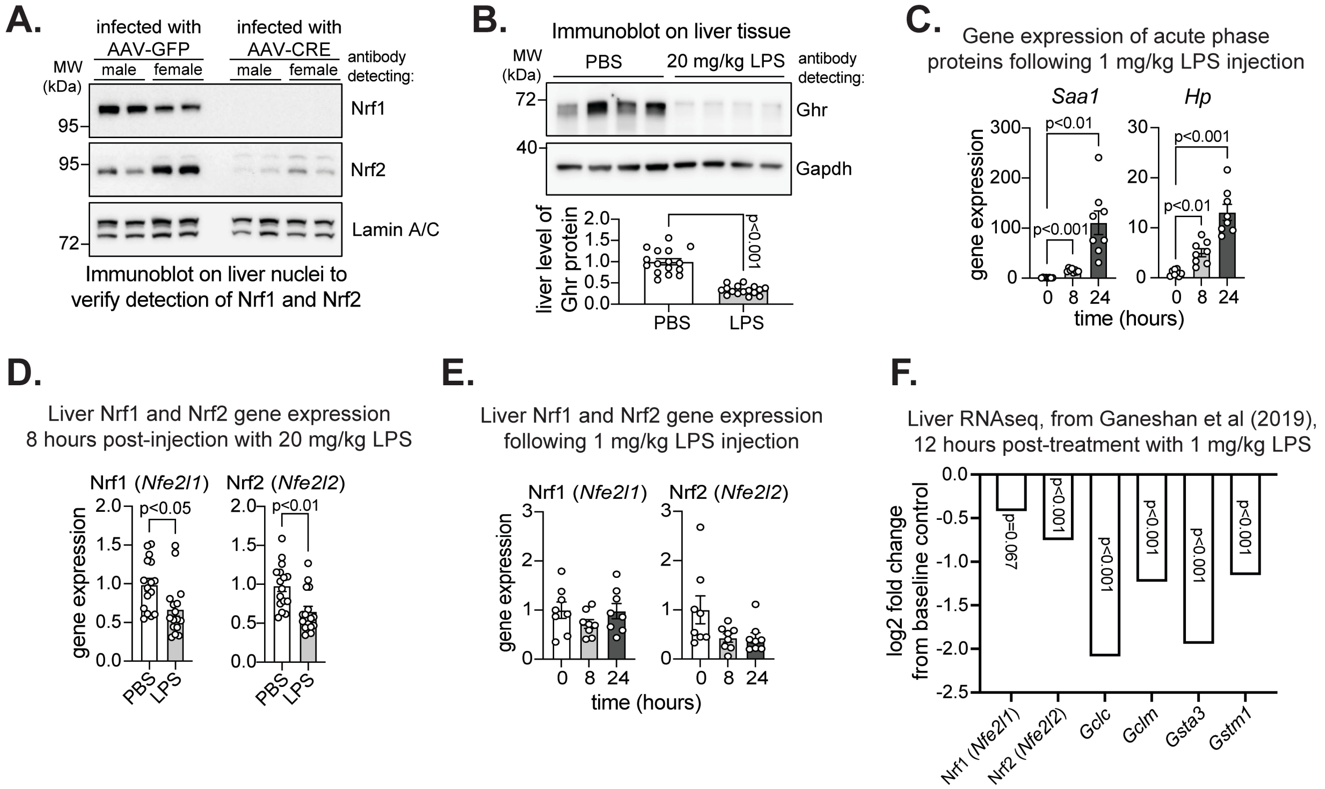


**Supplemental Figure 1. Effect of LPS on hepatic Nrf1 and Nrf2.** A) Immunoblot of liver nuclei from Nfe2l1flox/flox; Nfe2l2flox/flox mice infected with AAV-GFP or AAV-CRE to verify the molecular size of Nrf1 and Nrf2 (n = 2 males; 2 females). Immunoblot of lamin A/C was a loading control. B) Immunoblot and analysis of liver lysate to verify 20 mg/kg LPS reduced liver growth hormone receptor (Ghr), normalized by glyceraldehyde 3-phosphate dehydrogenase (Gapdh), which is expected to occur in sepsis (n = 8 males; 8 females). C) Liver expression of acute phase response genes 0-24 hours post-injection with 1 mg/kg LPS, normalized by 36b4 (n = 4 males; 4 females). D) Liver expression of Nrf1 (Nfe2l1) and Nrf2 (Nfe2l2) genes 8 hours post-injection with 20 mg/kg LPS or PBS, normalized by 36b4 (n = 8 males; 8 females). E) Liver expression of Nrf1 (Nfe2l1) and Nrf2 (Nfe2l2) genes 0-24 hours post-injection with 1 mg/kg LPS, normalized by 36b4 (n = 4 males; 4 females). F) Re-analyzed RNAseq data from Ganeshan et al. (2019), showing changes in Nrf1 (Nfe2l1) and Nrf2 (Nfe2l2) genes and their target genes 12 hours post-injection with 1 mg/kg LPS. p-value in B and D determined by t-test. p-value in C and E determined by one-way ANOVA, with Dunnett's post-test. p-value in F determined by Benjamin Hochberg analysis. Data in B-E are mean ± standard error of the mean, with individual data points shown.

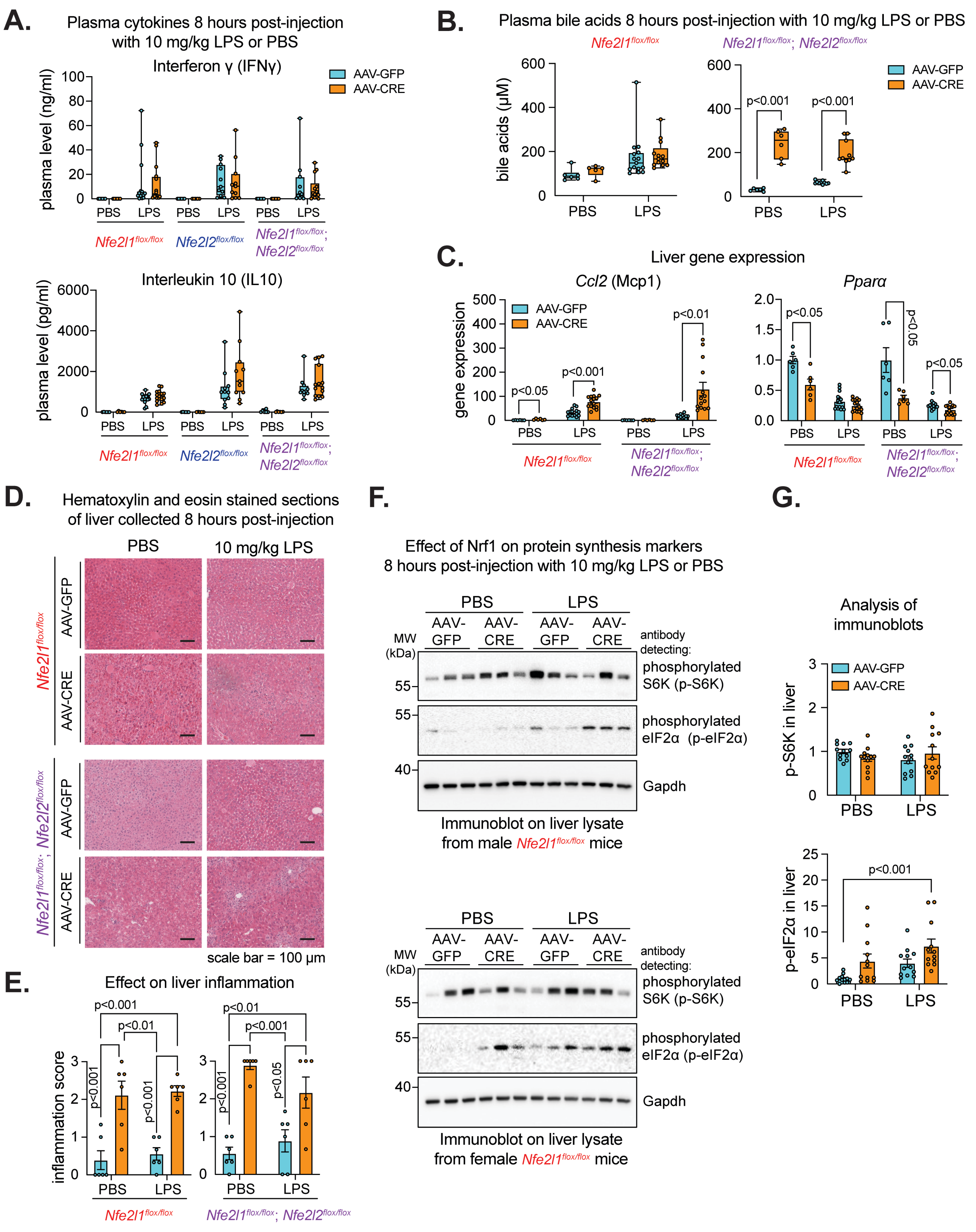


**Supplemental Figure 2. Effect of hepatic Nrf1, Nrf2, and combined deficiency on liver of mice with LPS-induced sepsis.** A-C) Plasma cytokines (A) and bile acids (B) as well as expression of indicated genes in liver, normalized by 36b4,(C) of indicated flox mice 8-hours post-injection with 10 mg/kg LPS or PBS (n = 3-7 males; 2-8 females). D-E) Hematoxylin and eosin-stained liver sections (D), with scale bar in panel, and corresponding liver inflammation score (E) of indicated flox mice 8-hours post-injection with 10 mg/kg LPS or PBS (n = 3 males; 3 females). F-G) Immunoblot of indicated phosphorylated protein in liver lysate (F) and its analysis (G), normalized by glyceraldehyde 3-phosphate dehydrogenase (Gapdh) (n = 6 males; 6 females). p-values in A and C determined by t-test, with adjustment for multiple comparisons. p-values in B, E, and G determined by two-way ANOVA with Tukey's post-test. Data in A-B are box and whisker plots, with individual data points shown. Data in C, E, and G are mean ± standard error of the mean, with individual data points shown.

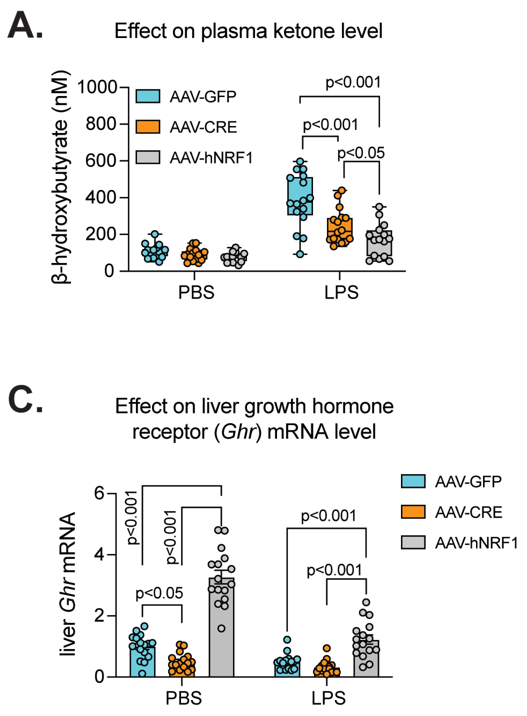


**Supplemental Figure 3. Effect of hepatic Nrf1 loss- and gain-of-function in mice with LPS-induced sepsis.** A-B) Plasma ketone β-hydroxybutyrate (A) and IGF1 (B) comparing control to Nfe2l1flox/flox mice with hepatic Nrf1 loss- or gain-of-function 8-hours post injection with 20 mg/kg LPS or PBS (n = 7-8 males; 8 females). C-D) Liver expression of genes encoding Ghr and Ccl2 8-hours post injection with 20 mg/kg LPS or PBS, normalized by 36b4 (n = 8 males; 8 females). p-values in A-D determined by two-way ANOVA with Tukey's post-test. Data in A-B are box and whisker plots, with individual data points shown. Data in C-D are mean ± standard error of the mean, with individual data points shown.

**A.**

Principal component analysis (PCA): liver RNAseq on LPS exposed *Nfe2l1*$^{flox/flox}$ mice infected with AAV-GFP or AAV-CRE

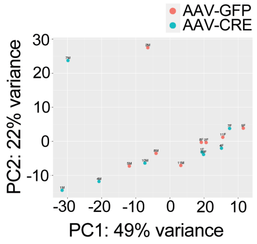


Number of differentially expressed genes:
AAV-CRE vs AAV-GFP = 0

**B.**

Principal component analysis (PCA): liver RNAseq on LPS exposed *Nfe2l1*$^{flox/flox}$ mice infected with AAV-GFP or AAV-hNRF1

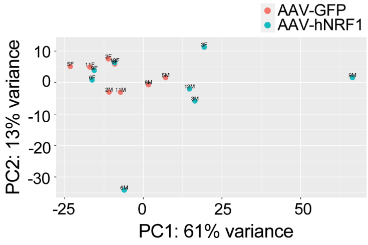


Number of differentially expressed genes:
AAV-hNRF1 vs AAV-GFP = 330

**Supplemental Figure 4. Dimensional analysis of RNAseq data.**
A-B) Principal component analysis for RNAseq profiles of liver from Nfe2l1flox/flox control mice (AAV-GFP) versus mice with hepatic Nrf1 loss-of-function (AAV-CRE) (A) or hepatic Nrf1 gain-of-function (AAV-hNRF1) (B) (n= 4 males; 4 females). Number of differentially expressed genes for each comparison are shown in bottom of each panel.

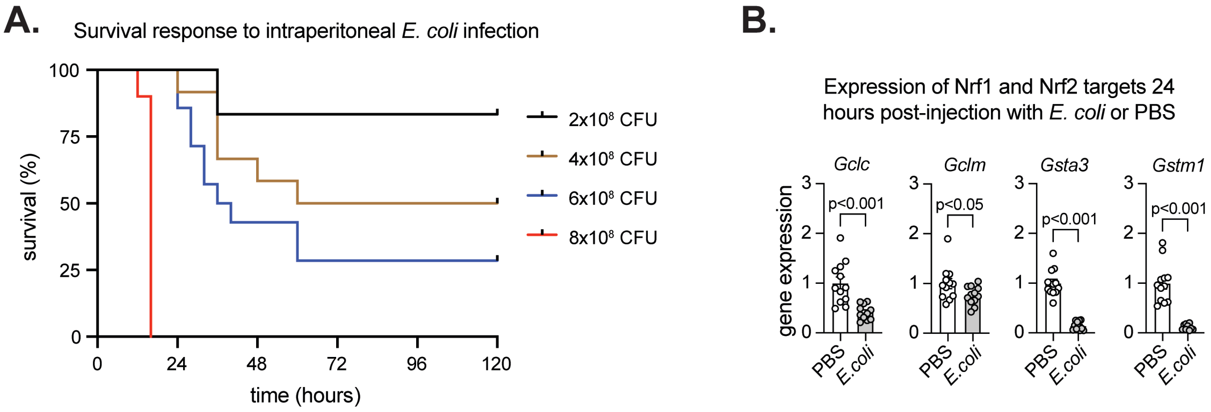


**Supplemental Figure 5. Establishment of an E. coli-induced sepsis model in the Nfe2l1flox/flox mice.**
A) Survival plot of Nfe2l1flox/flox mice intraperitoneally (IP) injected with indicated colony forming units (CFU) of E. coli (n = 5-6 males; 5-8 females). B) Liver expression of Nrf1 and Nrf2 target genes 24-hours post injection with PBS, 4 x 108 CFU of E. coli (males), or 3.2 x 108 CFU of E. coli (females), normalized by 36b4, (n= 4-6 males; 6 females). p-value in B determined by t-test. Data in B are mean ± standard error of the mean, with individual data points shown.

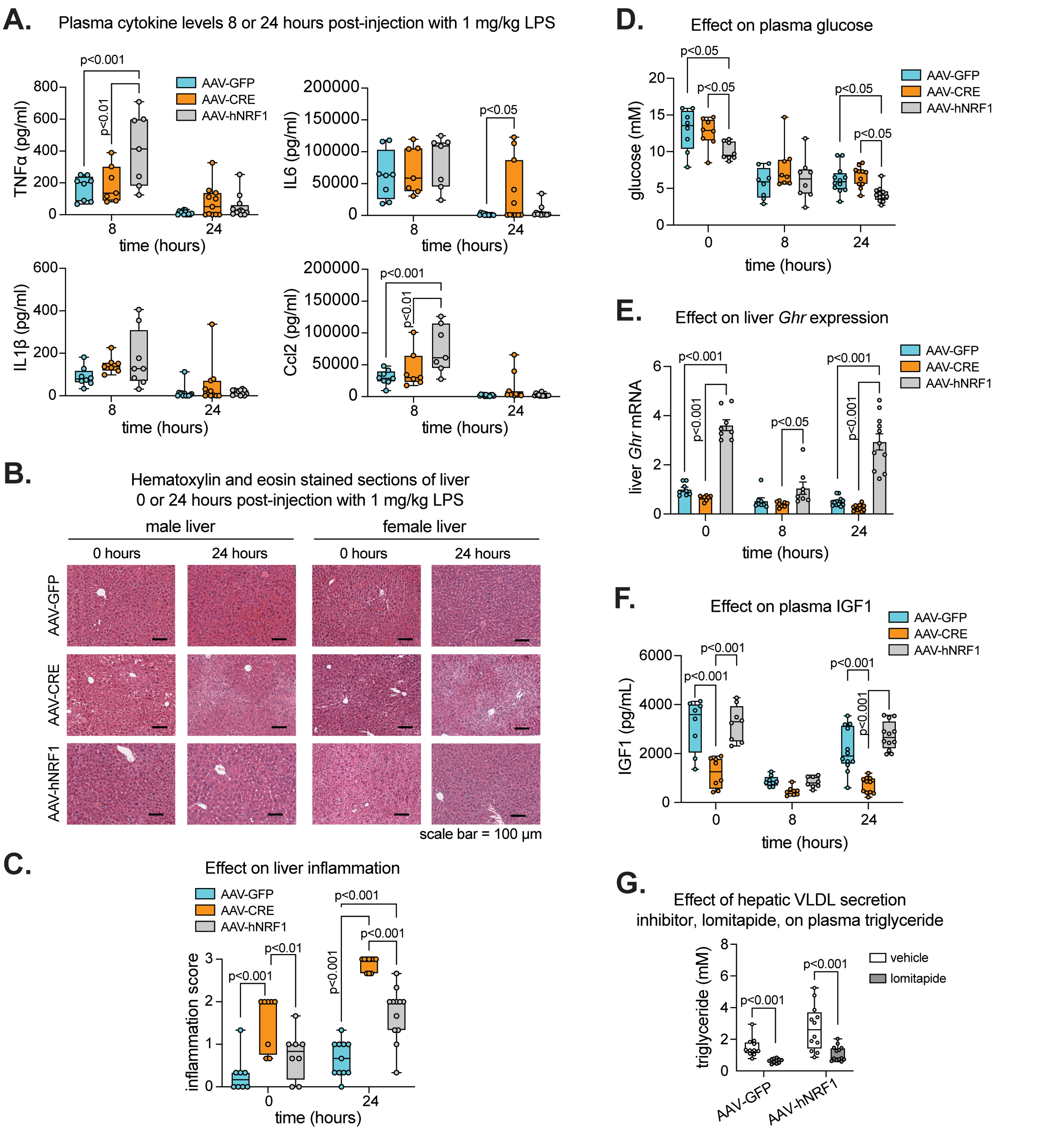


**Supplemental Figure 6. Effect of 1 mg/kg LPS on Nfe2l1flox/flox mice with hepatic Nrf1 loss- and gain-of-function.** A) Plasma cytokines, comparing control to Nfe2l1flox/flox mice with hepatic Nrf1 loss- or gain-of-function 0-24 hours post-injection of 1 mg/kg LPS (n = 3-6 males; 4-6 females). B-C) Hematoxylin and eosin-stained liver sections (B), with scale bar in panel, and corresponding liver inflammation scores (C) comparing control to Nfe2l1flox/flox mice with hepatic Nrf1 loss- or gain-of-function 0-24 hours post-injection of 1 mg/kg LPS (n = 4-6 males; 4-6 females). D-F) Plasma glycemia (D), liver expression of Ghr, normalized by 36b4, (E), and plasma IGF1 (F) comparing control to Nfe2l1flox/flox mice with hepatic Nrf1 loss- or gain-of-function 0-24 hours post-injection of 1 mg/kg LPS (n = 4-6 males; 4-6 females). G) Plasma triglyceride in Nfe2l1flox/flox control (AA-GFP) mice and mice with increased hepatic Nrf1 activity (AAV-hNRF1) injected with PBS or 2 mg/kg lomitapide to confirm reduced hepatic VLDL secretion (n = 5-6 males; 5-6 females). p-value in A, C-F determined by two-way ANOVA, with Tukey's post-test. p-value in G determined by t-test, with adjustment for multiple comparisons. Data in A, C-D, and F-G are box and whisker plots, with individual data points shown. Data in E are mean ± standard error of the mean, with individual data points shown.
