## Supplemental Table S4 for "Hepatic Nrf1 (*Nfe2l1*) promotes VLDL dependent liver defense against sepsis"

**Supplemental Table 2. List of primer sequences used in qPCR**

| **Gene** | **Forward primer** | **Reverse primer** |
|---|---|---|
| *36b4* | AGGGCGACCTGGAAGTCC | CCCACAATGAAGCATTTTGGA |
| *Ccl2 (Mcp1)* | GATCACCAGCAGCAGGTGT | ATGTATGTCTGGACCCATTCCTT |
| *Gclc* | AACAAGAAACATCCGGCATC | CGTAGCCTCGGTAAAATGGA |
| *Gclm* | TCCCATGCAGTGGAGAAGAT | AGCTGTGCAACTCCAAGGAC |
| *Ghr* | CTGCAAAGAATCAATCCAAGCC | CAGTTCAGGGGAACGACACTT |
| *Gsta3* | AAGAATGGAGCCTATCCGGTG | CCATCACTTCGTAACCTTGCC |
| *Gstm1* | ATACTGGGATACTGGAACGTCC | AGTCAGGGTTGTAACAGAGCAT |
| *Hp* | GCTATGTGGAGCACTTGGTTC | CACCCATTGCTTCTCGTCGTT |
| *Nfe2l1 (Nrf1)* | GACAAGATCATCAACCTGCCTGTAG | GCTCACTTCCTCCGGTCCTTTG |
| *Nfe2l2 (Nrf2)* | CAGCTCAAGGGCACAGTGC | GTGGCCCAAGTCTTGCTCC |
| *Ppara* | AGAGCCCCATCTGTCCTCTC | ACTGGTAGTCTGCAAAACCAAA |
| *Saa1* | TTGTTCACGAGGCTTTCC | TGAGCAGCATCATAGTTCC |
| *Tbp* | GAGCTGTGATGTGAAGTTTCC | TCTGGGTTTGATCATTCTGTAG |
