## Supplemental Table S5 for "Hepatic Nrf1 (*Nfe2l1*) promotes VLDL dependent liver defense against sepsis"

**Supplemental Table 3. Primary antibodies used for immunoblots**

| Antibody | Supplier | Catalog number; RRID |
|---|---|---|
| anti-Gapdh | Invitrogen | AM4300; AB_2536381 |
| anti-Ghr | Santa Cruz Biotechnologies | Sc-137185; AB_2111405 |
| anti-Lamin A/C | Cell Signaling Technologies | 4777S; AB_10545756 |
| anti-Nrf1 | Cell Signaling Technologies | 8052S; AB_11178947 |
| anti-Nrf2 | Cell Signaling Technologies | 12721S; AB_10891429 |
| anti-p65 (NF-κB) | Cell Signaling Technologies | 8242S; AB_10859369 |
| anti-p-eIF2α | Cell Signaling Technologies | 3398T; AB_2096481 |
| anti-p-S6K | Cell Signaling Technologies | 9234S; AB_2269803 |
| anti-Mouse HRP | Cell Signaling Technologies | 7076S; AB_330924 |
| anti-Rabbit HRP | Cell Signaling Technologies | 7074S; AB_2099233 |
